# TCRdenoise - an unsupervised similarity-based approach for denoising of TCR-pMHC specificity data

**DOI:** 10.64898/2026.09.03.749070

**Authors:** Johanne Margrethe Lund, Sebastian Nymann Deleuran, Morten Nielsen

## Abstract

Public repositories of T cell receptor (TCR)-peptide-MHC (pMHC) interactions constitute a critical resource for studying adaptive immunity and developing predictive models of TCR specificity. However, recent evidence suggests that a substantial fraction of reported TCR-pMHC interactions may be incorrectly annotated, limiting the quality of downstream analyses and machine learning applications. Here, we present an unsupervised sequence similarity-based framework for denoising peptide-specific TCR repertoires. The method combines pairwise TCR similarity metrics derived from TCRbase and TCRdist3 with hierarchical clustering and a novel adaptation of the silhouette score designed to address the prevalence of singleton clusters and highly imbalanced cluster structures. By incorporating a pseudo-cluster containing singleton and background TCRs, and by optimising both clustering distance thresholds and minimum cluster-size criteria, the proposed approach identifies TCRs likely to represent true antigen-specific binders while filtering putative noise.

Using experimentally validated repertoires from TCRvdb, we demonstrate that the modified silhouette score closely tracks clustering solutions that maximise separation between binding and non-binding TCRs, achieving strong agreement with independent validation based on the Matthews correlation coefficient. Extension to a large collection of peptide-specific TCR data revealed a strong negative correlation between the percentage of TCRs classified as noise and the predictive performance of peptide-specific binding models. Further, the denoising classification labels on this data set were corroborated using structural modeling confidence scores of the peptide-TCR interface extracted from a refined AlphaFold 3 modeling pipeline. Additionally, retraining NetTCR on denoised data improved internal cross-validated performance compared with models trained on the full data set, whereas models trained exclusively on TCRs classified as noise performed close to random.

Together, these results demonstrate that sequence similarity-based denoising can effectively enrich for biologically meaningful TCR-pMHC interactions and improve the quality of training data for predictive immunological models. The proposed framework provides a scalable strategy for improving the reliability of public TCR databases and facilitating the development of more accurate TCR specificity prediction methods.

## Introduction

A central component of the adaptive immune system is T cells. They recognise antigenic peptides presented by the major histocompatibility complex (MHC) on the surface of a host cell through their T cell receptor (TCR), forming the TCR-peptide-MHC (TCR-pMHC) complex [1]. The TCR consists of an α- and β-chain; each chain has a variable region containing three complementarity-determining regions (CDRs) that mediate binding to the pMHC [2]. Among these, the CDR3 regions exhibit the greatest sequence diversity and primarily interact with the peptide, whereas CDR1 and CDR2 interact mainly with the MHC molecule [1].

Although these databases are essential for computational immunology, they are known to contain a high level of label noise, meaning that an unknown proportion of the data may share incorrect pMHC annotations. For VDJdb, a recent study demonstrated that in the context of two peptides, approximately 50% of the reported TCR-pMHC could be experimentally validated binders [3]. Incorrect specificity annotations have large implications for TCR research. Training machine learning models, such as TCR binding prediction models, on noisy data will likely result in inferior predictive performance. In addition, noisy training data can obscure the ability of models to identify and learn underlying biological patterns [4]. Although experimental re-evaluation efforts, such as the validated VDJdb subset (TCRvdb) [3], improve annotation quality, these approaches are costly and cannot easily be expanded to keep pace with the rapid growth of repertoire sequencing data or the large volume of existing data requiring validation. Therefore, computational methods capable of identifying and removing likely non-binding TCRs are essential for improving the quality of publicly available data sets and enabling the development of more accurate predictive models.

Sequence similarity is a measure of homology between proteins, and homologous proteins often share similar functionality such as binding specificity [5]. Although T-cell receptors differ from most proteins in that their CDRs are generated through somatic V(D)J recombination rather than inherited sequence variation, TCRs that recognize the same pMHC frequently share conserved sequence features [3], [6]. This makes sequence similarity a meaningful measure for identifying TCRs with shared peptide specificity. The relationship between sequence similarity and peptide recognition also provides a basis for computational denoising of peptide-specific TCR repertoires. It is expected that within a repertoire, binding TCRs have greater sequence similarity to one another than non-binding TCRs [3]. The work by Messemaker et al. demonstrated the credibility of this approach, wherein they observed that validated binding TCRs can more frequently be grouped in clusters with high intra-similarity compared to non-validated binders [3]. However, a critical issue faced when using such similarity-based approaches for clustering and denoising is the challenge associated with the identification of the optimal clustering solution [7]. This in particular for data sets characterized by high cluster size imbalances, i.e., many singletons and/or a few very large clusters [8].

Here, we seek to address this issue and develop an unsupervised sequence similarity-based method to identify noise in TCR-pMHC repertoires from databases such as VDJdb. Using data from TCRvdb, NetTCR2.2, and IMMREP2023, we showcase the performance of the method for denoising both in terms of data set statistics and impact on training and evaluation of prediction models such as NetTCR.

## Methods

### Clustering evaluation data

The data set used in formulating TCRdenoise was obtained from the TCRvdb [3] and covered TCRs specific for two peptides, YLQPRTFLL (YLQ) and GLCTLVAML (GLC). Each data point contains the TCRs’ binding status (binder or non-binder). From the TCRvdb, 3 sequences from GLC and 19 from YLQ had no adjusted p-values and were removed.

Furthermore, when extracting the CDRs from the data, Stitchr failed to generate the full TCR sequence for 11 TCRs (1 from YLQ and 10 from GLC), resulting in further removal of these data points from the data set. Additionally, one of the binding TCRs for GLC was removed due to redundancy. Finally, the GLC data set contains 182 TCRs, with 131 binders and 51 non-binders. The YLQ data set contained 420 TCRs with 172 binders and 248 non-binders. All data were paired α- and β-chain TCR sequences.

### Background data

A TCR background data set was obtained from the IMMREP22 negative control data set [9], which contains 15,957 TCRs. The data set was downsized to 6368 TCRs using a Hobohm1 algorithm with a distance threshold of 0.3, with the input of a distance matrix from TCRdist3. These TCRs were then randomly downsampled to 1000 TCRs. This was done to ensure that the background TCRs represented a diverse range of sequences while minimizing sequence redundancy. The background TCR data set was included in the peptide-specific TCR repertoires as preparation for denoising as described later.

### TCR sequence similarities

Pairwise distances between TCRs were calculated using TCRbase and TCRdist3, which quantify sequence similarity by comparing the complementarity-determining regions (CDRs) of TCRs. In short, TCRbase uses K-mer similarity and BLOSUM62 to determine the sequence similarities, whereas TCRdist3 uses a distance-modified BLOSUM62 matrix to calculate the distance between TCRs. Both use weights for their calculation between the CDRs, where TCRbase was used with (1:1:4) and TCRdist3 (1:1:3) for CDR1, CDR2 and CDR3, respectively. TCRdist3 also has the option of including CDR2.5 as input. This was excluded from this study because the background data set lacked this information. The pairwise distance matrix was normalized to range between 0 and 1 and served as input to Agglomerative Clustering. Note that the background TCR data set was included in the pairwise distance calculations.

### Clustering

Hierarchical clustering was performed using agglomerative clustering with a single-linkage criterion. The input was the pairwise distance matrix (normalized to fall in the range 0-1) and 200 distance thresholds between 0 and 1. The distance thresholds were defined by selecting 200 logarithmically spaced quantiles spanning 0.0001 to 1 from the unique pairwise distances in the matrix to highlight how the data behaves during clustering. Before clustering, background TCRs were removed from the distance matrix to avoid distorting the clustering, and they were assigned to the same pseudo-cluster as the singletons (see below).

### Silhouette score

The Silhouette Score (SI_**i**_) evaluates the quality of a clustering solution by comparing the average distance between a TCR and members of its own cluster (a) with the average distance to the nearest neighboring cluster (b) (see equation 1). Scores range from −1 to 1, with higher values indicating better cluster separation. The global silhouette score (SI) was calculated as the mean SI_**i**_ score across all TCRs (see equation 2) [10].

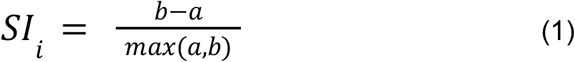

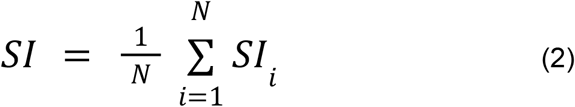

While the SI score is widely used for cluster validation, it is not directly applicable to denoising. This is due to singletons having no intracluster distance, and the SI score needing a minimum of two clusters to calculate the intercluster distance [10]; both of which limit the interpretability of the standard SI score for denoising. To address these limitations, we developed a modified SI score and singleton handling consisting of three components: a pseudo-cluster grouping singletons, inclusion of background TCRs in the pseudo-cluster, and a modification of the global SI score calculation.

The modified SI score works by calculating the SI_**i**_ score, similar to the default. However, the two methods differ in how they calculate the global SI score. Here, the modified version sums individual scores, including only the clustered TCRs (Nc) (see equation 3).

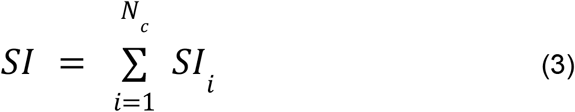

Since background TCRs are part of the pseudo-cluster, they function as a secondary cluster even when all the other TCRs form one cluster, enabling calculation of the SI score when all other TCRs are clustered together. Further, a search was incorporated to identify a size threshold for clusters to be included in the pseudo-cluster to allow filtering of small clusters of non-binding TCRs sharing similar sequence features. The search cumulatively reassigned clusters to the pseudo-cluster starting from the smallest cluster. At each step, the global SI score is calculated, and the threshold with the largest SI score is determined to be the solution.

For the distance threshold yielding the highest SI score, an additional search was performed to identify the minimum cluster size threshold that maintained an SI score at least 95% of the maximum. The smallest threshold was selected to minimize the effect of overfitting and limit the number of clusters incorporated into the pseudo-cluster while maintaining near-optimal SI performance.

### Validation

The Matthews Correlation Coefficient (MCC) was used as an independent metric to evaluate how well each clustering solution separated binding from non-binding TCRs. MCC was calculated from a confusion matrix comparing the experimentally annotated binding status (binding versus non-binding) with the clustering assignment (clustered versus singleton/noise). For evaluating the correlation between the SI score and MCC across distance thresholds, the Pearson Correlation Coefficient (PCC) was calculated between the two scores.

### NetTCR training data

The data used for retraining NetTCR was an updated version of the NetTCR-2.2 training data set [11], which included combined entries from IEDB [12], VDJdb [13], and a 10x Genomics data set, resulting in 12,647 positive data points. In short, the NetTCR data set was constructed from CDR3s, with annotated V and J genes. Full-length TCRs were generated using Stitchr [14], and CDR region annotated using ANARCI [15]. The data set includes peptides with corresponding paired α- and β-chain sequences of the 6 CDRs. Negative data points were generated from the positive data by pairing each peptide with TCRs from peptides having a Levenshtein distance greater than 3 from the original peptide. Five negative examples were generated for each positive TCR, except for GILGFVFTL (1:4.42). Lastly, the data set was redundancy reduced and randomly partitioned into five subsets, and swapping was restricted to occur within each partition (for details on the data preparation pipeline, refer to the original NetTCR-2.2 publication [11]).

From the positive data points, two other NetTCR training data sets were generated. These were generated based on the results from TCRdenoise. From the original 12,647 TCRs, 7518 were classified as binders and 5,129 were classified as noise. These data sets were further subsampled to ensure a minimum of five positive TCRs per partition and the same peptide composition, resulting in 5,910 in the denoised training data and 4,585 data points in the noise training data covering the peptides GLC, GIL, IVT, NLV, RAK, TPR, YLQ, and YVL.

### IMMREP2023 Data

The IMMREP2023 data set [2] covered both private and public data, totaling 3484 data points. The data set contains positive and negative TCRs but was a subset containing data points from the peptides GLC, GIL, IVT, NLV, RAK, TPR, YLQ, and YVL, the same peptides that the NetTCR models are trained on (see above).

### Training of NetTCR

NetTCR is an ensemble of convolutional neural networks used for predicting TCR-pMHC binding [11]. Each input sequence is passed through a CNN layer, and the max-pooled output is fed into a dense layer and onward to the output neuron (for details, refer to Jensen and Nielsen [11]). In this study, we retrain NetTCR on the full, denoised, and noise NetTCR training data described above. The models were trained in a 5-fold nested cross-validation setup. For each model, the performance evaluation was done in a per-peptide manner in terms of the AUC0.1 value calculated from the concatenated test set predictions and on unseen data from IMMREP2023. For details on the model training, model architecture, and performance evaluation, refer to the original NetTCR-2.2 publication [11].

### NetTCRfold

NetTCRfold is an AlphaFold 3-based pipeline adapted for TCR-pMHC structural modeling. In short, the adaptation consists of tailoring the sequence databases used for MSA generation and template search to only include TCR-pMHC-relevant data, a query-limited approach for template identification, and a refined confidence metric, ipSAE d0dom, predicting the confidence of the peptide-TCR interface used for model selection and model scoring. For further details, refer to [16].

## Results

To improve the separation of antigen-binding and non-binding T cell receptors (TCRs) within immune repertoires, we here developed a sequence-based denoising framework that leverages TCR similarity within a given data set. The method first computes pairwise distance matrices using both TCRbase and TCRdist3. For each distance matrix, a series of agglomerative clustering solutions is generated across a range of distance thresholds. Next, we develop a refined SI score modified for handling singleton clusters to determine the optimal clustering threshold. Finally, we investigate how the optimal clustering outputs derived from the complementary distance measures can be integrated into a consensus clustering approach, providing enhanced discrimination between binding and non-binding TCR populations. Finally, we assess the performance of this framework and its ability to enrich antigen-specific TCRs while reducing repertoire noise in a set of benchmark conditions.

### Clustering

We first illustrate the development of the denoising pipeline in the context of the TCRvdb data. The first step consists of the construction of pairwise TCR distance matrices. Figure 1 shows these matrices for the TCRs associated with the two peptides (YLQ and GLC) in the TCRvdb data, calculated for TCRbase and TCRdist3, respectively. From this figure, it is apparent that TCRs for the two peptides cluster quite differently. YLQ appears to have one main cluster with high purity of annotated binders, and the non-binding TCRs have overall high mutual distances, meaning low similarity with other TCRs, indicating that a good separation between binding and non-binding TCRs is possible. In contrast, GLC has multiple disconnected clusters, suggesting that the separation between binding and non-binding TCRs is more difficult.

**Figure 1:**
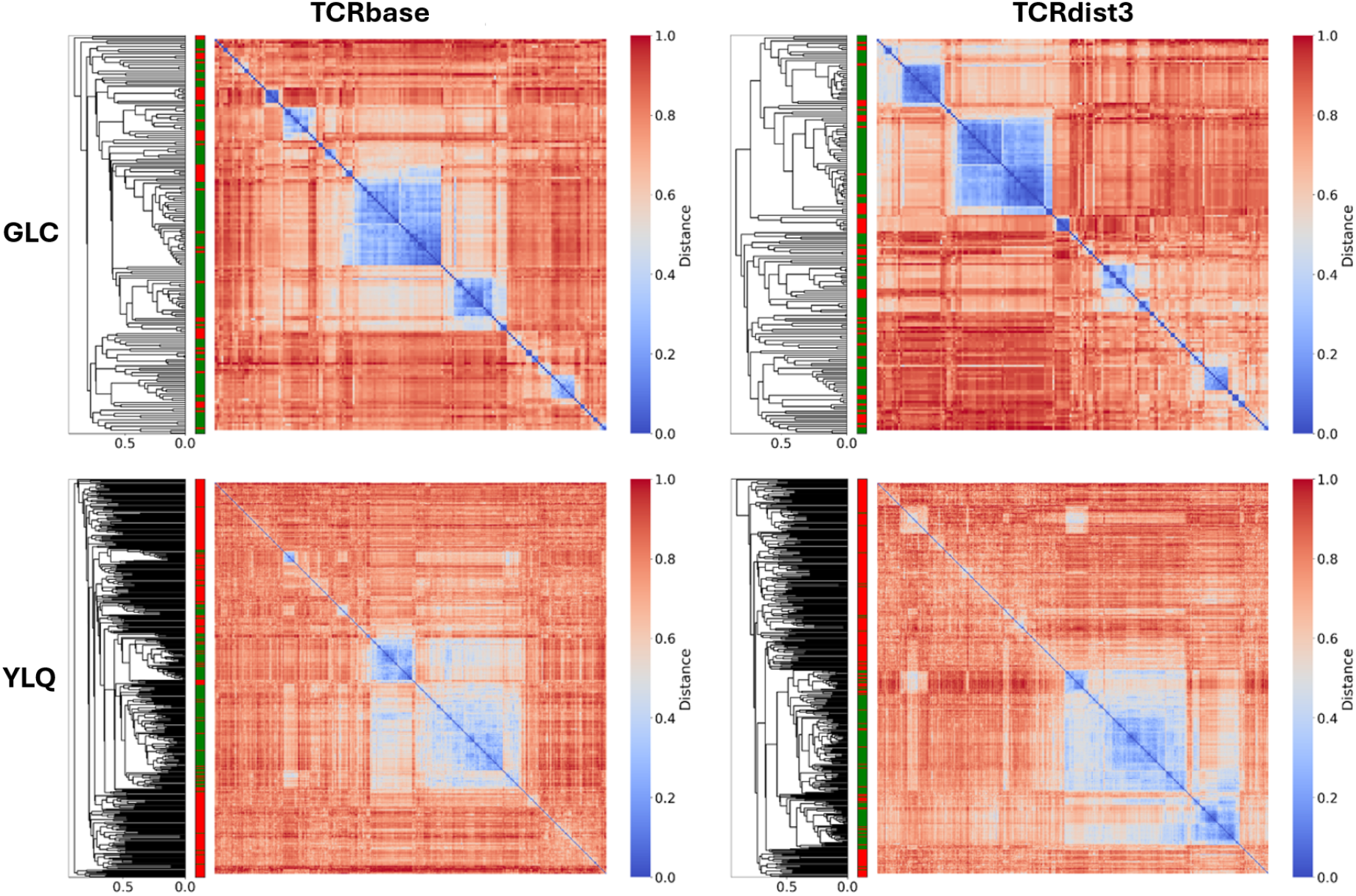
Distance Matrix Heatmap and Dendrogram from TCRbase and TCRdist3 for the TCRvdb data sets for the peptides GLC and YLQ. The distance matrix was sorted by the dendrogram generated from Agglomerative clustering used on the distance matrices computed from the two methods. The color block on the left side of the heatmaps indicates binding status for each TCR: non-binder (red) and binder (green). For GLC, there are 182 TCRs (131 binders, 51 non-binders), and for YLQ, 420 TCRs (172 binders, 248 non-binders). Each distance matrix was normalized to fall in the range 0-1.

### Default SI score

To evaluate the clustering solutions generated by the clustering, we first investigated the use of SI score as a validation metric. To determine whether the distance threshold resulting in the highest SI score also provided the best separation between binding and non-binding TCRs, the Matthews correlation coefficient (MCC) was calculated for the clustering solution at each distance threshold. The MCC was calculated from a confusion matrix comparing the annotated TCR binding status (binding versus non-binding) with the clustering assignment (singleton versus clustered). Next, curves were generated showing the SI and MCC values as a function of the clustering distance threshold (see Figure 2).

**Figure 2:**
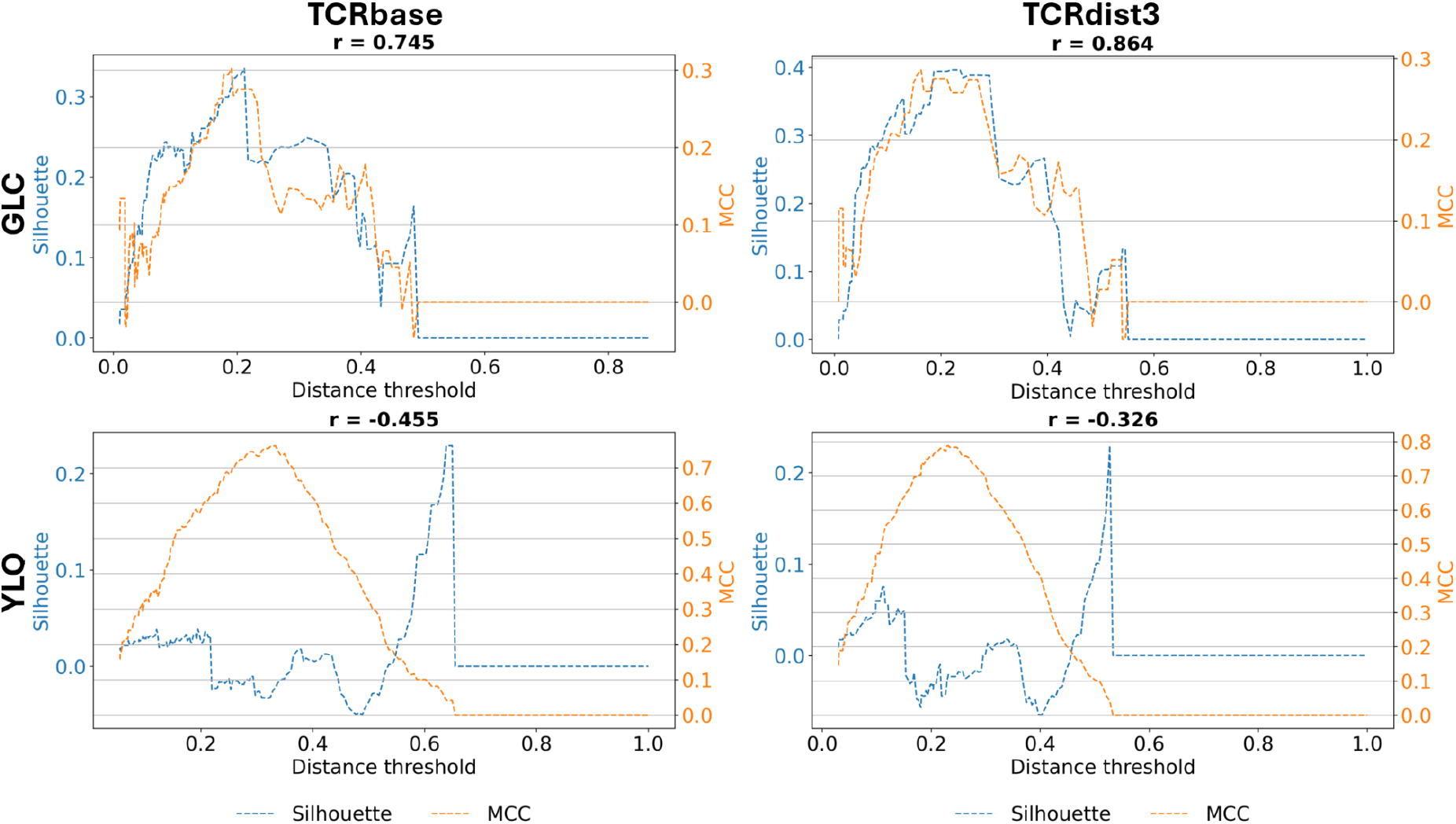
Relation between clustering threshold and SI and MCC values using the default SI metric. The x-axis represents the distance threshold. The blue y-axis (left) and curve represent the SI score, and the orange y-axis (right) and curve the MCC. SI score and MCC were generated based on the clustering solutions generated by Agglomerative clustering. Clustered TCRs were classified as binders, and singletons were classified as noise. The SI score used is the default by Scikit-Learn (Equations 1 and 2). MCC was calculated for each clustering solution, where the clustered TCRs were classified as binders, and the singletons were classified as noise. Each subplot’s title includes the PCC’s r-value for correlation between the SI curve and MCC curve.

The results of this analysis revealed a strong correlation between the MCC and SI curves for GCL, which was further supported by high positive PCC values. However, for YLQ, the same high and positive correlation was not observed. These results contrasted at first with our initial expectations of which of the repertoires would be easier to denoise, based on the clustering observed in Figure 1.

### Redefining the SI score

The clustering initially assigns each TCR to its own cluster and subsequently merges clusters as the distance threshold increases. As a result, singleton clusters are present across many distance thresholds. Since intra-cluster distance cannot be calculated for singletons, their SI value is undefined. To account for this limitation, singletons were grouped into a pseudo-cluster that was excluded from the global SI score calculation. Consequently, the global score was calculated as the sum rather than the mean of the remaining cluster SI scores, thereby avoiding normalization over a varying number of contributing clusters.

Although this modification reduced the influence of singleton clusters, a second limitation became apparent at larger distance thresholds. As clusters continued to merge, the singleton pseudo-cluster could eventually contain only one or a few TCRs, while the remaining data formed a single large cluster. This not only makes the SI score sensitive to the composition of the pseudo-cluster but also prevents its calculation once only a single cluster remains. To overcome these limitations, a set of background TCRs was incorporated into the pseudo-cluster, ensuring that it remained sufficiently large across all distance thresholds and allowing SI scores to be calculated throughout the clustering process (for details on the background TCR data set, refer to methods).

The results of these updates to the SI score calculation are shown in Figure 3. After incorporating the background data and modifying singleton handling, a small decrease in PCC was observed for GLC. However, the performance was greatly increased for YLQ. While this performance increase was limited in terms of the PCC value, the maxima of the SI curves now corresponded more closely to the maxima of the MCC curves across all four data sets (see Figure 3), indicating that the modified SI score more reliably identified the clustering solutions that best separated binding and non-binding TCRs.

**Figure 3:**
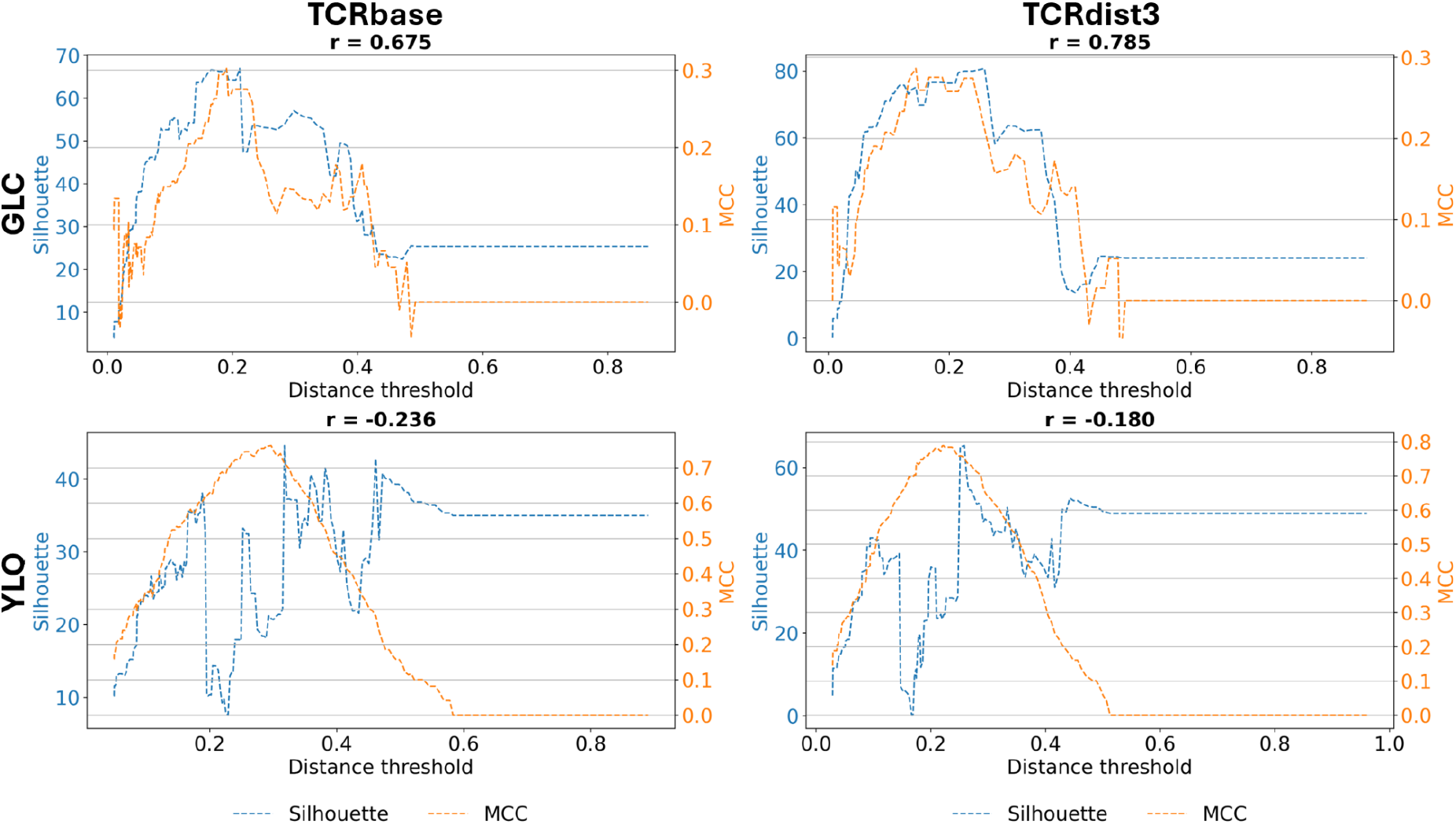
Relation between clustering threshold and SI and MCC values using the SI metric modified to include the singleton pseudo-cluster. The x-axis represents the distance threshold. The Blue y-axis (left) and curve are the SI score, and the orange y-axis (right) and curve are the MCC. The data sets had an additional 1000 background TCRs functioning as a permanent part of the pseudo-cluster. SI score was calculated as the sum of each clustered TCR’s SI score (Equation 3). MCC was calculated for each clustering solution, in which the clustered TCRs were classified as binders and the pseudo-cluster as noise. Each subplot’s title includes the PCC’s r-value for correlation between the SI curve and MCC curve.

### Optimal size threshold

Despite this improvement, the SI curve for YLQ still displayed several local maxima and minima, which were not observed for GLC, suggesting that, at least for YLQ, small non-singleton clusters could represent noise. To address this, a parameter search was performed over all possible cluster size thresholds, where clusters with sizes equal to or below the selected threshold were assigned to the pseudo-cluster. At each threshold, the modified SI score was calculated, and the size threshold yielding the highest score was selected (for details on this cluster size search, refer to methods).

After implementing the cluster size threshold search approach into the SI score calculation, a clear correlation between MCC and SI score was observed across all four data sets (see Figure 4). Furthermore, the previously observed peaks and valleys in the YLQ SI curves were no longer present, indicating improved agreement between the SI-based validation and the biological binding annotations.

**Figure 4:**
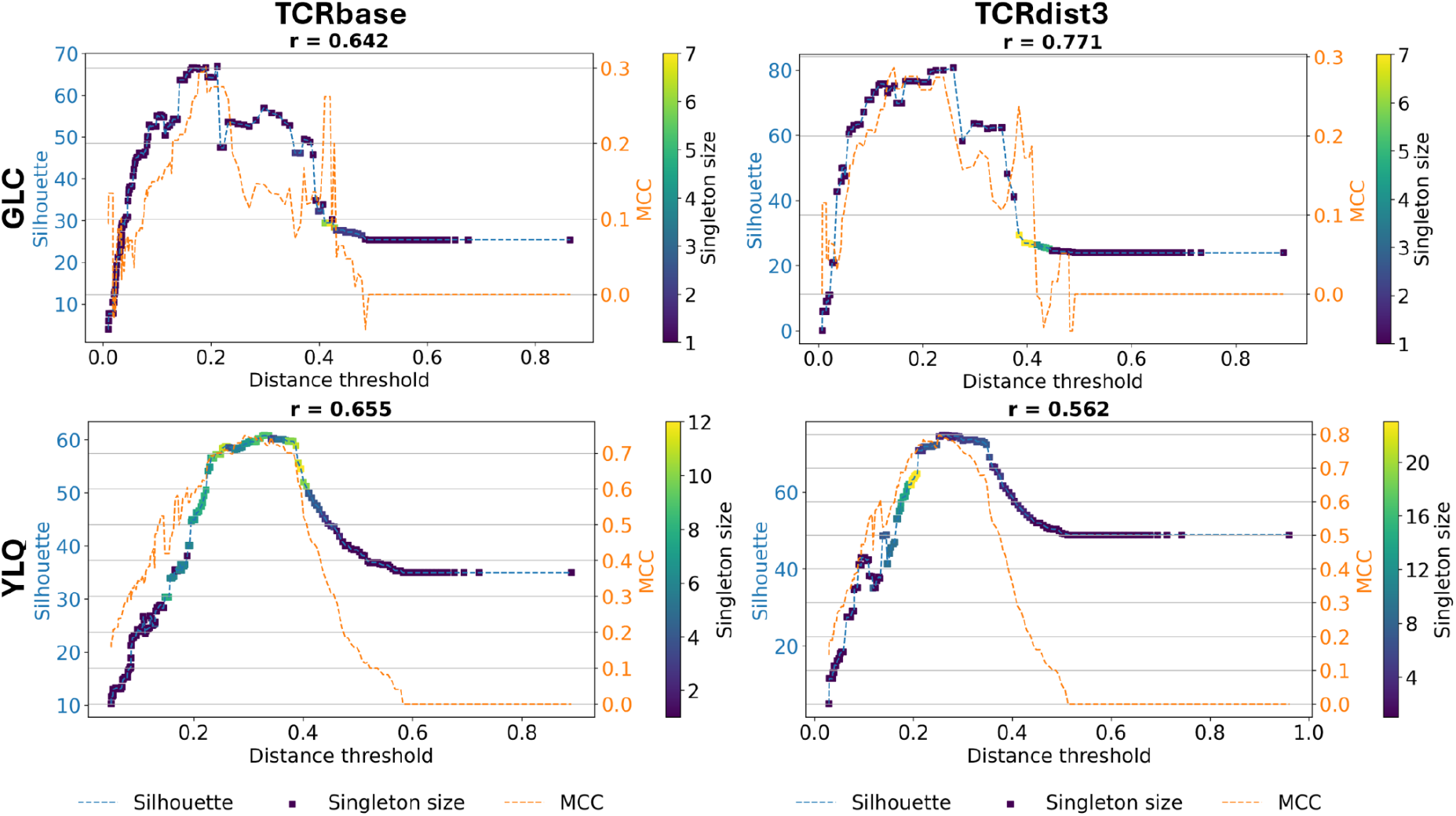
Relation between clustering threshold and SI and MCC values using the SI metric modified to include the singleton pseudo-cluster and cluster size threshold optimization. The x-axis represents the distance threshold. The blue y-axis (left) is the SI score, and the orange y-axis (right) is the SI-specific MCC. The coloration of the plot points on the SI curve depicts the different pseudo-cluster size thresholds (Size threshold). The SI score was calculated as the sum of each clustered TCR’s SI score (Equation 3). At each distance threshold, the SI score was calculated for different size thresholds. The highest SI score was then chosen for this distance threshold. The size thresholds determine at what size clusters are grouped with the pseudo-cluster. MCC was calculated for the highest SI scores, in which the clustered TCRs were classified as binders and the pseudo-cluster as noise. Each subplot’s title includes the PCC’s r-value for correlation between the SI curve and MCC curve.

### Consensus

After determining the clustering solution for each similarity method (TCRbase and TCRdist3), a consensus solution was created from the union of clustered TCRs in the individual cluster solutions. While this consensus strategy slightly reduces overall mean accuracy, it serves as a meaningful compromise between the two individual solutions, for both peptides achieving an improved or comparable performance compared to the lowest-performing method. Furthermore, adopting the union of both methods maximizes downstream data retention, filtering out only those data points flagged as noisy by both methods (see Table 1).

**Table 1:** MCC for the different methods and the consensus between them.

| Method | GLC - MCC | YLQ - MCC | Average MCC |
| --- | --- | --- | --- |
| Consensus | 0.212 | 0.746 | 0.479 |
| TCRbase | 0.275 | 0.729 | 0.523 |
| TCRdist3 | 0.212 | 0.779 | 0.535 |

### In silico validation

Up to this point, method development and evaluation have been limited to the two peptides in the TCRvdb data set. To further validate TCRdenoise on a broader data set, we next used the positive data points from the NetTCR training data for in silico validation (for details on this data set, refer to methods). This data set does not include labels of validated binders or non-binding “noise”. Rather, we compared properties of data sets associated with high versus low proportions of “noise”, and shared characteristics of TCRs classified as binders versus noise in terms of NetTCR prediction and AlphaFold 3 structural modeling scores.

First, we retrained NetTCR on the full NetTCR data set and reported the cross-validated predictive performance (for details on the model trained, refer to methods). Next, we plotted the proportion of denoised TCRs as a function of the NetTCR predictive performance for each peptide. The resulting scatterplot (Figure 5A) revealed a strong negative correlation (PCC = -0.69) between the percentage of TCRs classified as noise and the AUC0.1. The proportion of noise identified tended to correlate with data set size. That is, for data sets with similar AUC0.1 values, the peptides with a high number of data points shared a higher proportion of identified noise, particularly for the subset of peptides with AUC0.1 values less than 0.8, as perceived by the color of the dots in Figure 5A.

**Figure 5:**
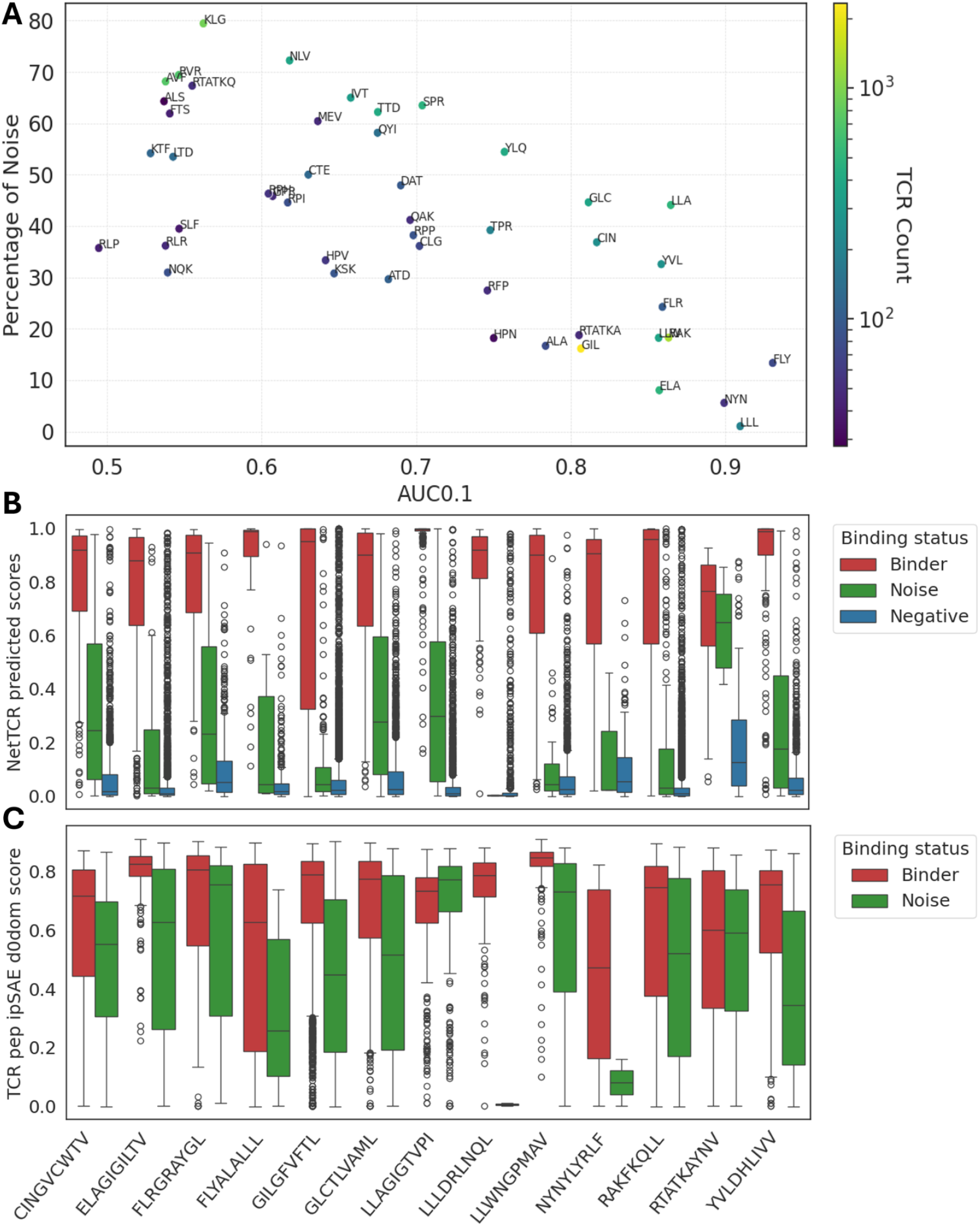
An investigation of the relations between the data separation and TCR-pMHC prediction scores. A) A scatter plot with NetTCR AUC0.1 on the x-axis and the percentage of noise found by TCRdenoise on the y-axis. NetTCR AUC0.1 values were obtained in cross-validation. Each point on the scatter plot is colored by the number of positive data points in the repertoire and has been labeled by its peptide name. B) The NetTCR prediction score distribution of the binders (red), the noise (green), and the swapped negatives (blue) for peptide repertoires with an AUC0.1 of 0.8 or greater. C) The distribution of TCR-pep ipSAE d0dom scores from NetTCRfold [16] for the predicted binders (red) and noise (green) for the same peptide repertoires as in panel B.

Next, we compared the distribution of NetTCR prediction scores for positives in the NetTCR data set classified as binders, as noise, and the swapped negatives, respectively, for the subset of peptides sharing a NetTCR cross-validated AUC performance above 0.8 (see Figure 5B). This figure clearly demonstrates that TCRs classified as noise by TCRdenoise overall share a higher prediction score overlap with the negative training data compared to the TCRs classified as binders. In concrete terms, for only one (RTATKAYNV) of the 12 peptides do the 25/75% percentile values for the binder and noise score distributions overlap. In contrast, this number is 10 out of 12 at the same percentile scores for the noise and negative score distributions (all peptides overlap except LLAGIGTVPI and RTATKAYNV). To understand why these outliers behave differently, we investigated the clustering solutions found in the two cases (Supplementary Figure 1). Here, we observe for LLAGIGTVPI that the clustering solutions are defined by one dominant, highly homogeneous cluster for both TCRbase and TCRdist3. TCRs in this large cluster share very high intra-similarity values, resulting in all the smaller, more heterogeneous clusters being assigned to the pseudo-cluster in the process of optimizing the SI score. The RTATKAYNV peptide is characterized by relatively few TCRs, and clustering solutions are defined by multiple clusters of varying size and intra-cluster similarity, while many clusters, in particular for the TRCbase solution, have a size of two to three. When looking at the NetTCR score distribution (Figure 5B), many of the TCRs labeled as noise appear to share scores comparable to the TCRs labeled as binders. This suggests that for this peptide, some of the singleton TCRs could be true binders to RTATKAYNV, underlining the key challenge of cluster-based denoising imposed by data size limitations.

Finally, we applied NetTCRfold, an AlphaFold 3-based structural modeling pipeline refined to predict TCR-pMHC specificity [16], to extract structural differences between TCRs labeled as binders versus noise. Here, structural models of each TCR-pMHC complex were generated using NetTCRfold, and the ipSAE model confidence score was calculated for each structure (for details on the NetTCRfold pipeline, refer to methods and Ballesteros-Cuartero et al. [16]). We next plotted the distribution of these ipSAE confidence scores for the TCRs labeled as binding and noise, respectively, for each of the peptides included in Figure 5B. Inspecting the result of this analysis, we observe the same tendency as seen in Figure 5B. In the vast majority of cases, the distribution of confidence scores for TCRs labeled as noise shares lower median values compared to the TCRs classified as binders. The only two exceptions to this are LLAGIGTVPI and RTATKAYNV, the same outliers as in Figure 5B.

Together, the results support the notion that the TCRs identified as noise by TCRdenoise to a very high degree are non-binders, but also, in the case of the two outlier peptides, point toward certain limitations of the method imposed mainly by data scarcity and the presence of highly imbalanced cluster size distributions.

### Retraining NetTCR

To further support the validity of the clustering results, we retrained NetTCR on different subsets of the NetTCR training data, namely the full, denoised, and noise data sets. Here, we reported both the internal cross-validated AUC0.1 performance for each pMHC across the different data sets and the model’s performance on the IMMREP23 data set (for details on model training, evaluation, and the IMMREP23 data set, refer to methods).

The result of this evaluation is shown in Figure 6. When considering the cross-validated NetTCR performance (Figure 6A), training on the denoised data set resulted, as expected, in higher performance compared to training on the full data set. Likewise, the performance on the noise data set was found to be very close to random. For the noise model, a similar behavior was observed for the IMMREP23 data (Figure 6B). Also, here the performance was close to random. In contrast, the two model trainings on the full and denoised data here demonstrated comparable performance. This latter observation is in alignment with the general observation that NetTCR is resilient to noise in its training data [11], and that such noise in many cases can serve as a source for model training regularization [17]. In conclusion, these results thus show that TCRs labeled as noise by TCRdenoise share no learnable signal and hence strongly support the conclusion that the clustering classification identifies TCRs not specific to their annotated pMHC.

**Figure 6:**
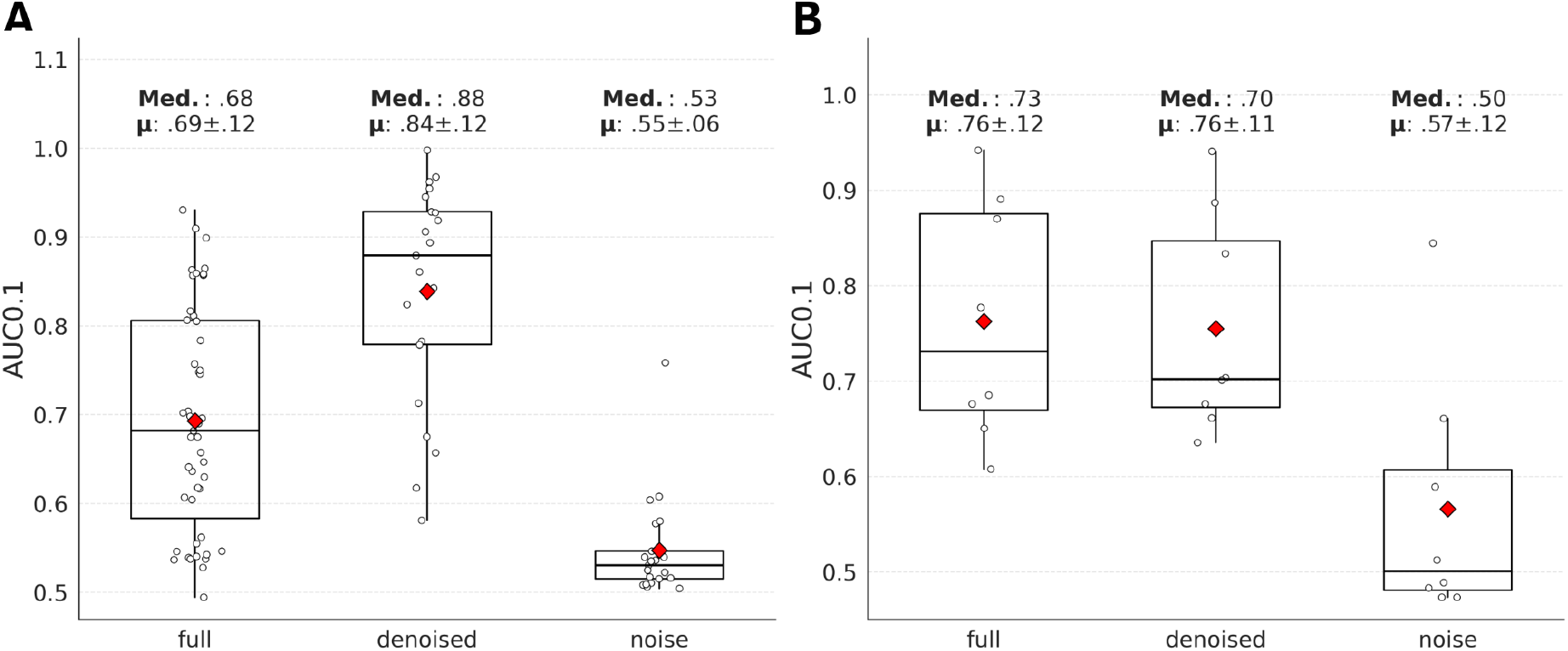
Boxplots with the AUC01 scores for the NetTCR models trained on either the full data set, the denoised data set, or the noise from the data set. A) Illustrates the models’ nested cross-evaluation on their respective training data. B) Illustrates the three models’ performance on a subset of the IMMREP2023 data set of both public and private data points limited to the peptide subset shared by all training data.

**Figure 7:**
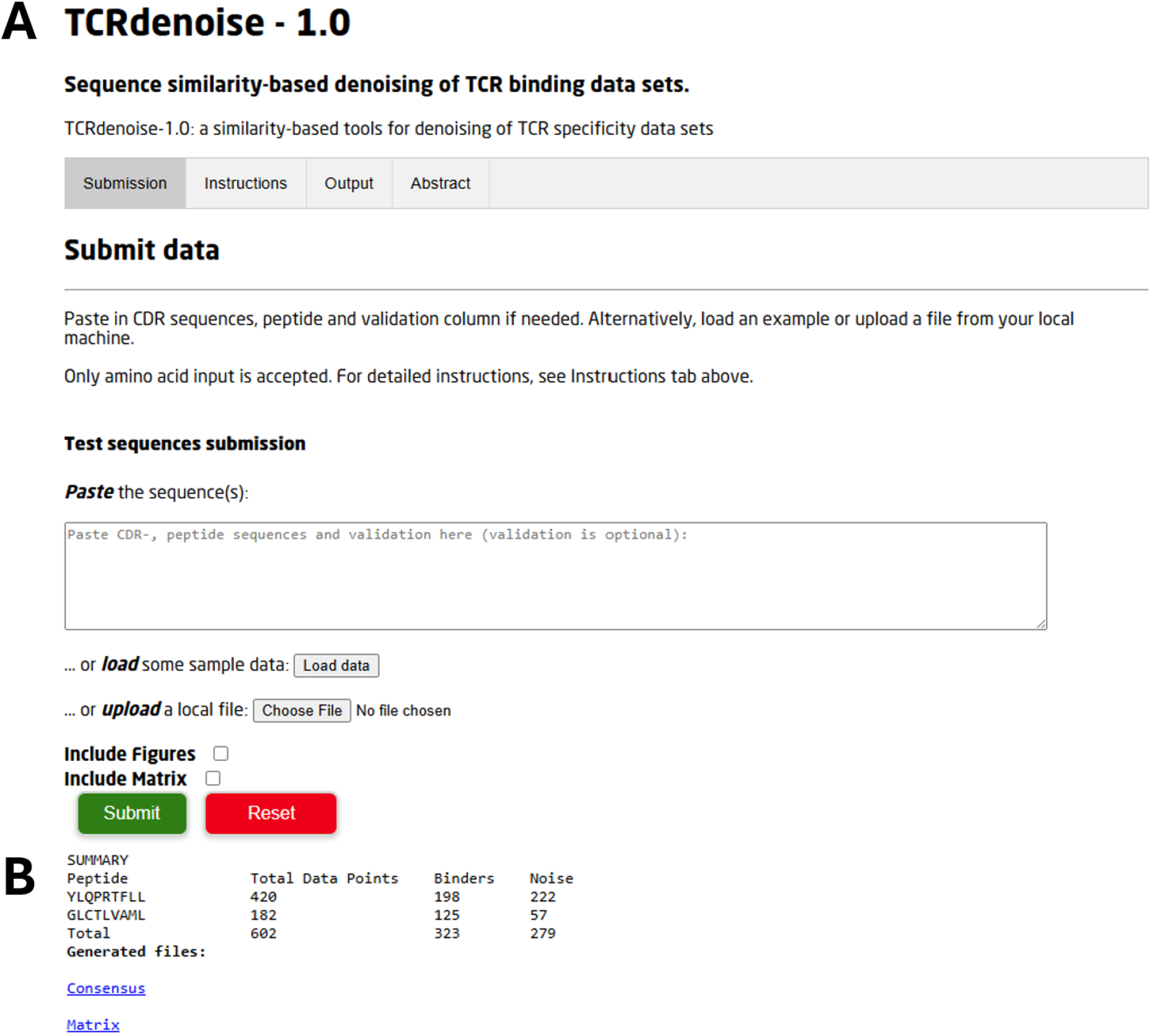
An overview of the web server. A) Illustrates the input section of the web server. B) shows the output when running the TCRvdb data set through the web server.

### The TCRdenoise method

The TCRdenoise method is implemented as a web server at https://services.healthtech.dtu.dk/services/TCRdenoise-1.0/. The method takes as input a data file with TCRs labeled to peptides and returns the TCRdist3, TCRbase, and consensus denoising solutions. For details, refer to the information on the website.

## Discussion

Here, we have developed TCRdenoise, a similarity-based method for denoising TCR-pMHC binding data sets and demonstrated its performance on both experimentally validated data sets and in the context of training refined in-silico prediction methods.

The characteristics of the TCRvdb data set used to develop and evaluate the TCRdenoise differed considerably between peptides. Whereas the YLQ repertoire consisted of a relatively homogeneous group of binding TCRs with low pairwise distances, the GLC repertoire displayed substantially greater sequence diversity among binders. These observations suggest that peptide-specific TCR repertoires can exhibit distinct similarity-distance distributions for binding TCRs, indicating the need for denoising methods that can adapt to these distributions rather than relying on fixed thresholds.

A key component of the proposed method relates to the refined definition of the silhouette metric and score used to define the optimal clustering solution. Here, we have demonstrated that simple updates, mainly related to the handling of singletons and inclusion of a negative set of background TCRs, address the main issues encountered when applying the default silhouette score for defining the optimal cluster solution.

Applying TCRdenoise to the TCRvdb data, we find an almost perfect association with the identified optimal cluster solution and the annotated TCR binding labels. This conclusion was extended to a large set of TCR-pMHC specificities from the public domain, where TCRs labeled as noise by TCRdenoise demonstrated strong shared evidence for binding non-binders using both machine learning and AlphaFold 3 structural model evaluation.

The limited amount of validated training data raises the possibility of overfitting. Design choices may unintentionally become tailored to the characteristics of the TCRvdb data sets rather than reflecting general properties of peptide-specific TCR repertoires. To address this, several components of TCRdenoise were made data-driven rather than fixed, including the dynamic distance threshold, the SI-based clustering solution, and the “singleton” size used to define the pseudo-cluster. Nevertheless, validation on additional experimentally verified data sets is necessary to further assess the robustness of TCRdenoise.

A critical limitation of TCRdenoise is its underlying assumption that binding TCRs form groups of sequence-similar receptors. While this assumption is supported by observations from TCRvdb, it is not necessarily universally valid for every binding TCR. This could result in validated binding TCRs with unique or highly divergent sequences being incorrectly classified as noise. This issue is clearly most critical in situations where the set of positive TCRs for a given peptide is sparse or when the TCR space is characterized by a few large dominant, highly homogeneous clusters. Note also that NetTCR to some degree suffers from the same limitation of mainly capturing signals from dominant homogeneous clusters during its training, while being challenged with capturing signals from singletons and small clusters. These observations make the underlying binder signal detection by TCRdenoise and NetTCR somewhat related and highlight that some caution should be taken when interpreting NetTCR predictions as independent validations of the TCRdenoise results.

However, the main finding in our analysis, namely that TCRs identified as noise share overlap with non-binding TCRs both in terms of NetTCR prediction scores and AF-3.0 structural modeling confidence values, still holds true, independent of these cautions.

From our results, we observed that for a few of the peptides where the TCR specificity space is characterized by a large, highly homogeneous cluster, other small minority clusters are removed from the optimal clustering solution. This occurs due to the dominant cluster having a low intracluster distance, resulting in a high SI score, leading to the size threshold search disfavoring solutions including the smaller, more heterogeneous cluster, as seen, for instance, for the poor-performing LLAGIGTVPI peptide. Another outlier in our analysis was RTATKAYNV. Here, the poor performance of TCRdenoise is mainly driven by the scarce amount of data points and low mutual similarity, resulting in solutions with many small clusters. This highlights the limitation that, in particular for small data sets, not all TCR singletons are necessarily noise.

While these remarks point to clear limitations of the proposed method, the fact that these are outlier observations suggests that the TCRdenoise in the vast majority of cases is robust, and only in the few borderline cases shares the risk of “over” denoising. This said, additional experimental validation will be essential to evaluate performance independent of sequence-similarity assumptions.

In conclusion, our results demonstrate that sequence similarity-based denoising can effectively distinguish signal from noise in peptide-specific TCR repertoires and improve the reliability of downstream computational analyses. Despite limitations related to repertoire diversity and data sparsity, the method shows robust performance across multiple independent validation strategies, supporting its utility as a practical preprocessing step for public TCR databases and future TCR specificity prediction frameworks.

## Funding / Acknowledgement

Research reported in this publication was supported by the Novo Nordisk Foundation Data Science Collaborative Research program under the grant NNF24OC0089619.

