## Supplementary Figure 1 for "TCRdenoise - an unsupervised similarity-based approach for denoising of TCR-pMHC specificity data"

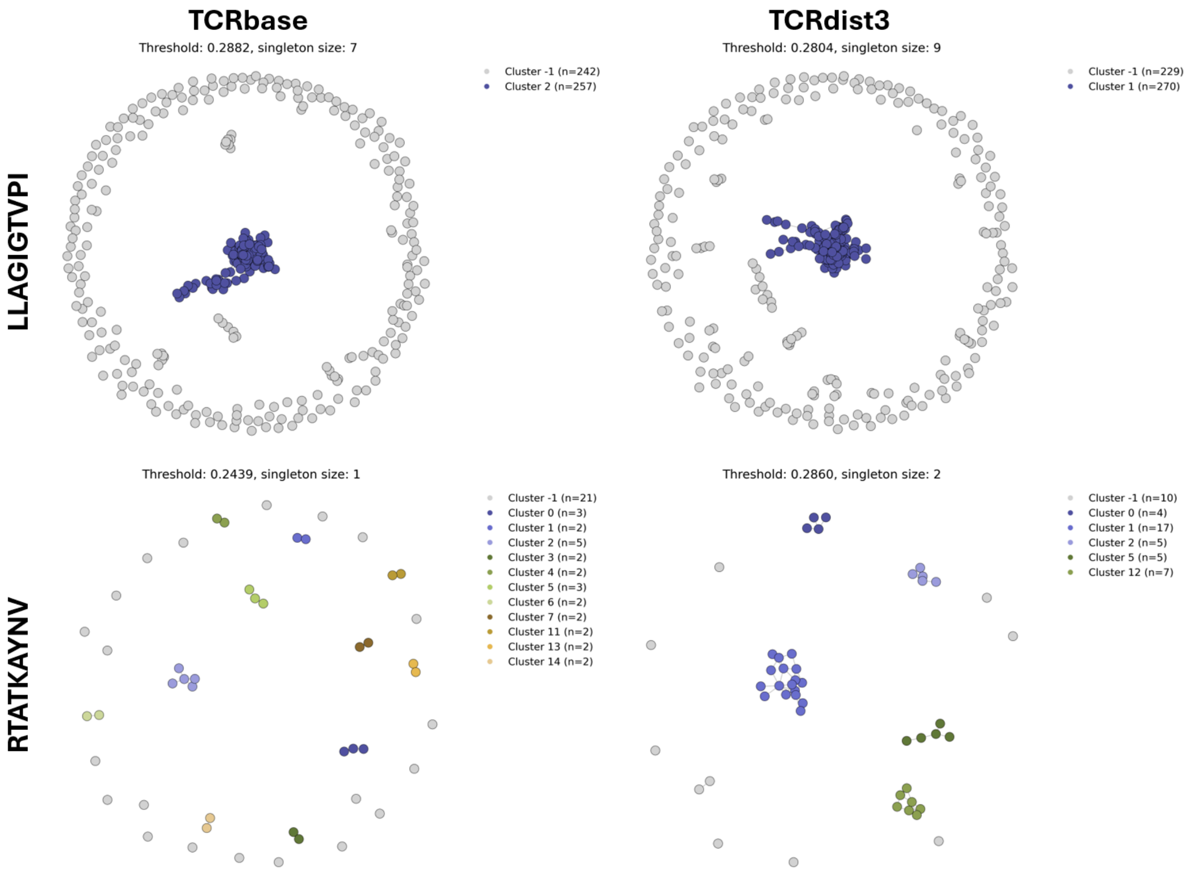


Supplementary Figure 1: Clustering solution of peptide LLAGIGTVPI and RTATKAYNV. The grey cluster -1 illustrates the data points found in the pseudo-cluster, excluding the background data. The colored data points are the clusters that TCRdenoise deems possible binders. Each subplot's title reports at what distance threshold the solution was formed and the “singleton” size threshold.

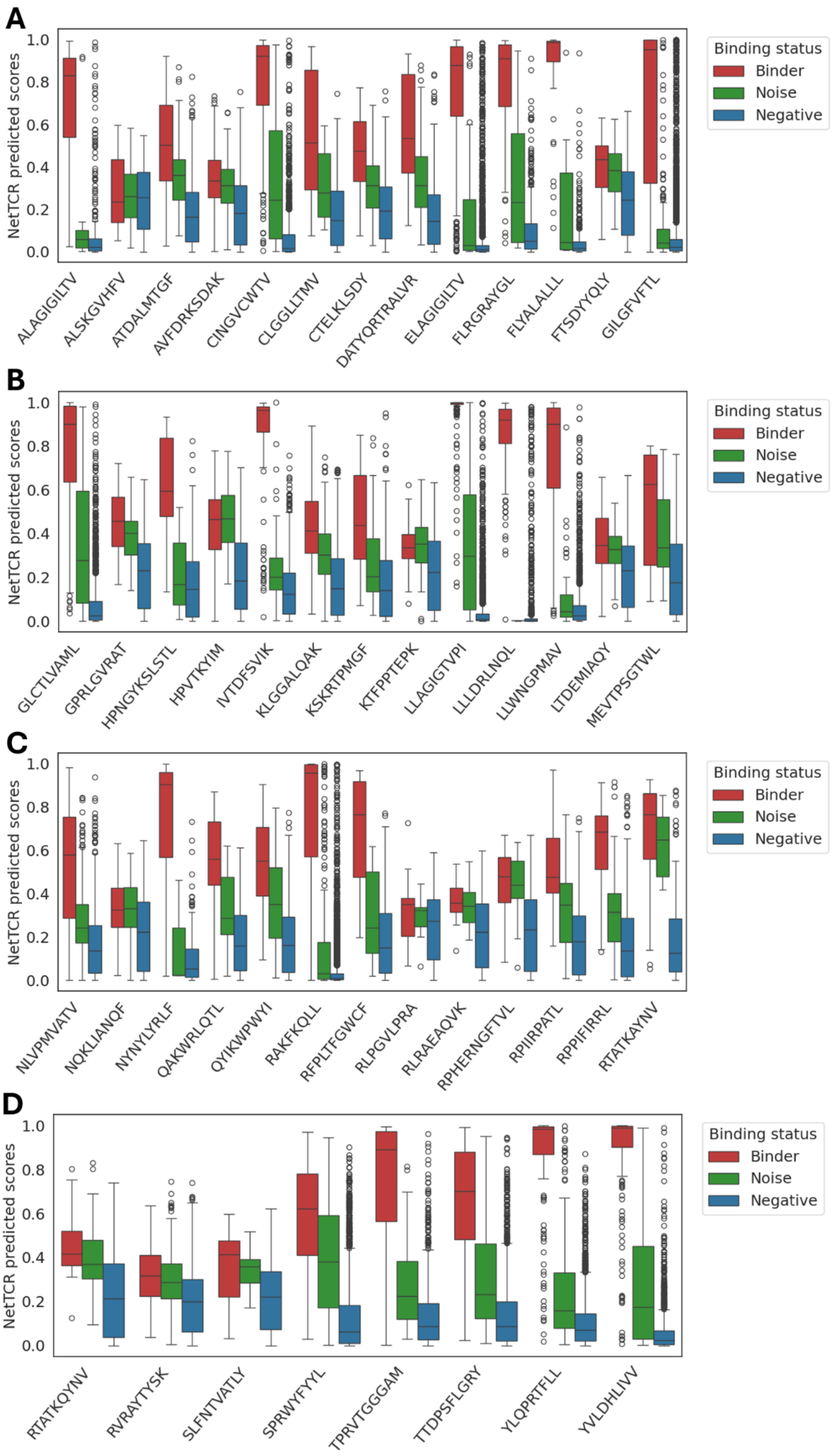


*Supplementary Figure 2: The NetTCR prediction score distribution of the binders (red), the noise (green), and the swapped negatives (blue).*

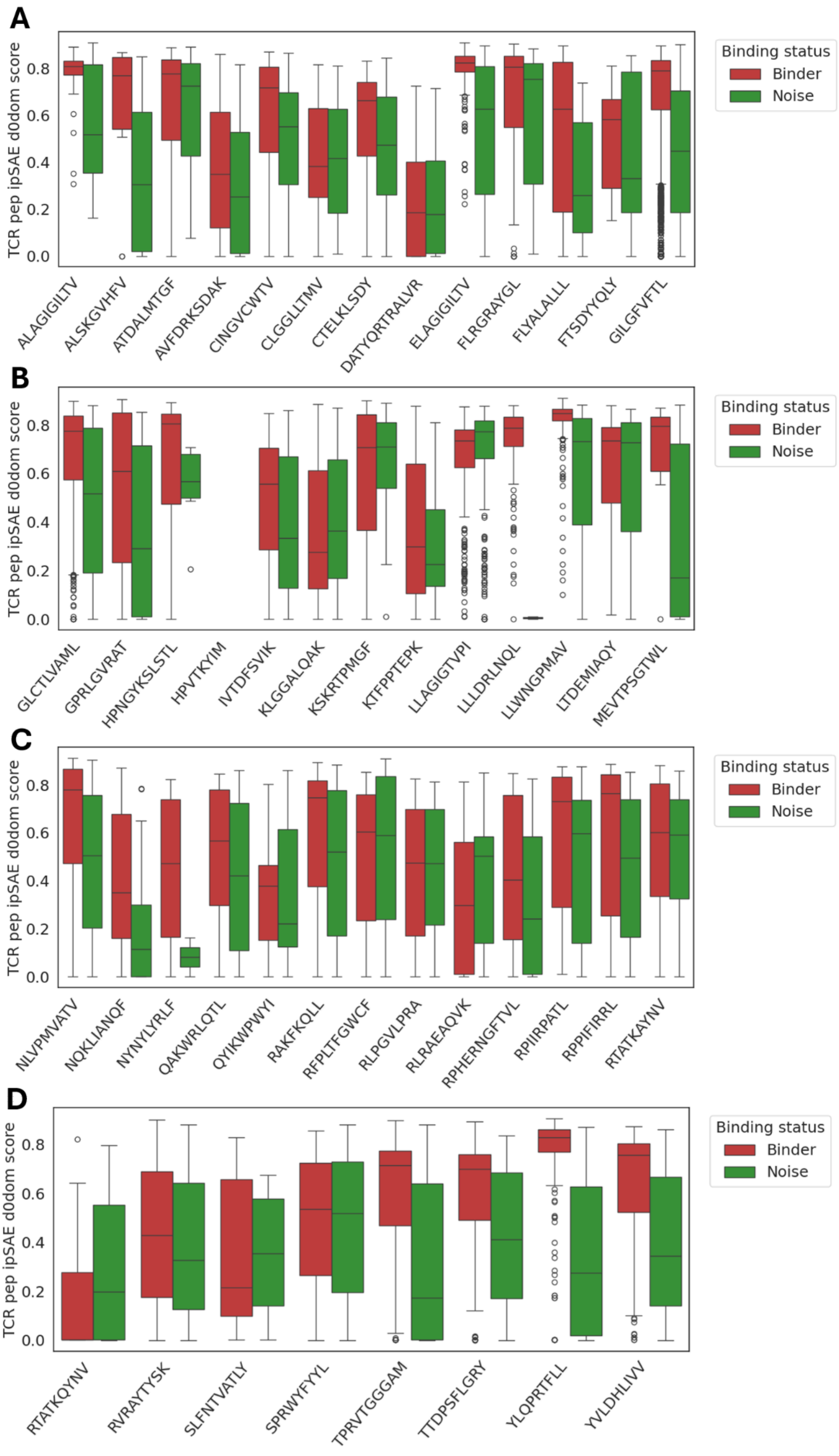


*Supplementary Figure 3: The distribution of TCR-pep ipSAE d0dom scores from NetTCRfold for the predicted binders (red) and noise (green).*
